# A high-throughput genomic screen identifies a role for the plasmid-borne Type II secretion system of *Escherichia* coli O157:H7 (Sakai) in plant-microbe interactions

**DOI:** 10.1101/2020.04.13.038984

**Authors:** Ashleigh Holmes, Leighton Pritchard, Peter Hedley, Jenny Morris, Sean P. McAteer, David L. Gally, Nicola J. Holden

## Abstract

Food-borne illness arising from Shiga-toxigenic *Escherichia coli* (STEC) is often linked to consumption of fruit and vegetables as the bacteria have the ability to interact with plants and use them as alternative or secondary hosts. The initial stages of the interaction involve chemotaxis, attachment and potentially, responding to the early stages of microbe perception by the plant host. We used a high-throughput positive-selection approach to identify early interaction factors of *E. coli* O157:H7 isolate Sakai to spinach. A bacterial artificial chromosome (BAC) clone library was quantified by microarray hybridisation, and gene loci enrichment measured using a Bayesian hierarchical model. The screen of four successive rounds of short-term (2 hour) interaction with spinach roots produced in 115 CDS credible candidates, comprising seven contiguous genomic regions. Two candidate regions were selected for functional assessment: a chaperone-usher fimbrial gene cluster (*loc6*) and the pO157 plasmid-encoded type two secretion system (T2SS). Interaction of bacteria with spinach tissue was reduced in the absence of the pO157 plasmid, which was appeared to involve the T2SS EtpD secretin protein, whereas loss of *loc6* did not impact interactions. The T2SS genes, *etpD* and *etpC*, were expressed at a plant-relevant temperature of 18 °C, and *etpD* expressed *in planta* by *E. coli* Sakai on spinach plants. Thus, a whole genome screening approach using a combination of computational modelling and functional assays has identified a novel function for STEC T2SS in interactions with plant tissue.

## 1 Introduction

Shiga-toxigenic *Escherichia coli* (STEC) (or verocytotoxigenic *E. coli*, VTEC) including the predominant serotype O157:H7, are significant zoonotic and food-borne pathogens, across the globe. Although ruminant farm animals are the primary reservoir for STEC, they can be transmitted through the food-chain on edible plants and plant-derived foodstuffs account for a large proportion (>50%) of food-borne illness in the USA [1]. However, animals remain the primary source of STEC on plants, either through direct application of manure/biosolids as fertilisers, or more likely via contaminated irrigation water [2].

STEC has been shown to interact with plants and can colonise them as secondary hosts [3]. Colonisation of STEC has been demonstrated on plant roots and in the rhizosphere [4–6], a favourable environment for bacteria that is rich in root exudates, which include a source of nutrients [7] and chemoattractants [8]. Numbers of *E. coli* recovered from roots often are greater than that from the leaves [6] and STEC has been shown to persist in soil and on plants for extended periods, e.g. >75 days [9].

Initial interactions in host colonisation involve chemotaxis, adherence and response to host perception. Since attachment is considered a prerequisite for successful colonisation, various approaches have been taken to identify adherence factors. The genome of STEC serotype O157:H7 isolate Sakai [10] encodes up to 14 fimbriae gene loci. Many of the *E. coli* adhesins show specificity in their host interactions, conferring a degree of tissue tropism for different *E. coli* pathotypes [11]. Curli, long polar fimbriae (Lpf), *Escherichia coli* common pilus (ECP), flagella and the T3SS have all been implicated in plant associated adherence of STEC [12–17], but several others STEC adherence gene clusters have yet to be functionally characterised. As such, we hypothesised that the STEC genome encodes additional uncharacterised factors that facilitate initial interactions with plant tissue. To identify which STEC genomic regions confer an advantage to colonisation of plant roots, a positive-selection screening approach was taken using an *E. coli* Sakai BAC clone library for short-term (2 hours) interactions with plant roots. Spinach was selected as it is relevant to large-scale STEC outbreaks [18], and we have previously shown specific adherence to spinach roots [14, 15, 6]. High-throughput screening enables wholesale analysis and previous global transcriptomic analysis has shown induction of STEC fimbrial and afimbrial adhesins in lettuce leaf lysates [19–21]. In a similar manner, high-throughput negative- and positive-selection approaches have identified colonisation factors, e.g. a random mutant library of *Pseudomonas fluorescens* was used to identify plant colonisation factors [22], and signature tagged mutagenesis and a bacterial artificial chromosome (BAC) library have been used to investigate STEC interactions with bovine mucus [23, 24]. Therefore, we used a BAC clone library of *E. coli* O157:H7 isolate Sakai that was previously used to identify genetic loci that enhanced adherence to bovine epithelial cells, and promoted bacterial growth in bovine mucus [23]. *E. coli* Sakai was used because it was derived from a large outbreak arising from contamination of white radish sprouts [25]. The approach involved whole-genome interrogation using microarrays and Bayesian analysis to compare the library clones prior- and post-spinach root inoculation.

The BAC library screen identified several contiguous *E. coli* Sakai chromosomal and plasmid regions that enriched following interaction with spinach roots, present on *E. coli* O157:H7-specific genomic segments known as S-loops [26]. Candidate regions that included annotated adherence factors were taken forward for characterisation. Functional analysis identified the plasmid-borne Type II Secretion System (T2SS) as a factor that conferred increased adherence for *E. coli* O157:H7 Sakai to both spinach roots and leaves.

## 2 Materials and Methods

### 2.1 Bacterial strains and media

*E. coli* O157:H7 isolate Sakai, hereafter *E. coli* Sakai [10] and its derivatives were grown in either lysogeny broth (LB) or MOPS medium [27] supplemented with 0.2 % glucose (or glycerol where indicated), 10 μM thiamine and MEM essential and non-essential amino acids (Sigma M5550 and M7145) termed rich defined MOPS (RD-MOPS) media. Antibiotics were included where necessary to maintain transformed plasmids at the following concentrations: 50 μg/ml kanamycin (Kan), 25 μg/ml chloramphenicol (Cam), 10 μg/ml Tetracycline (Tet), 50 μg/ml ampicillin (Amp).

### 2.2 Plant propagation

Spinach (*Spinacia oleracea*) cultivar Amazon seeds (Sutton Seeds, UK) were grown in hydroponics for the BAC screen. Seeds were germinated on distilled water agar (0.5 % w/v) and after 3-5 days transplanted into pots containing autoclaved vermiculite and sterile 0.5 × Murashige and Skoog (MS) medium (Sigma Aldrich, USA) with no carbon supplement. Plants were maintained under environmental cabinet conditions as above for 4-6 weeks. Spinach was grown similarly for BAC clone adherence assays and confocal microscopy of roots, for hydroponics plants in sterile hydroponic tubs (Greiner, UK) containing perlite instead of vermiculite (optimal for microscopy of roots). Spinach was grown in compost for adherence assays and confocal microscopy of leaves. Seedlings were grown in an environmental cabinet with a light intensity of 150 μmol m^2^ s ^−1^ (16 hour photoperiod) for a further 21 days at 22 °C. Compost-grown plants were germinated and maintained in individual posts with commercial compost and under glasshouse conditions 22 °C (16 h of light, 8 h of dark) with 130–150 μmol m^2^ s ^−1^ light intensity and 40% humidity.

### 2.3 Bacterial Artificial Chromosome Library screen for adherence to spinach roots

The BAC library contained a partial *Hind*III digest of *E. coli* Sakai genome cloned into pV41 vector, and together with the spinach root adherence approach, is described in detail in (Accompanying DiB paper DIB-S-20-00975).

### 2.4 Microarray hybridisation and data analysis

The microarray chip used for the analysis, a 8 × 15k *E. coli* gene expression array, *E. coli* v.2 (Agilent product number G4813A-020097) and *g*DNA extraction is described in detail in (*Submitted dataset to DiB*). Gene enrichment data is deposited with ArrayExpress with accession numbers for the adherence treatment: E-MTAB-5923 and control treatment: E-MTAB-5924. A complete description of the data analysis is provided at https://widdowquinn.github.io/SI_Holmes_etal_2017/ (doi:10.5281/zenodo.822825) but briefly, probe intensity data was subjected to QA and clean-up in which three problematic probes in a single treatment arm replicate were replaced with values interpolated from the other two treatment replicates. Array intensities were quantile normalised separately for control and treatment arms, and each probe annotated by BLASTN match to the most recent CDS annotations for the *E. coli* DH10B and Sakai isolates (NCBI accessions: GCF_000019425.1_ASM1942v1, GCF_000008865.1_ASM886v1). Only probes that unambiguously matched to a single Sakai or DH10B CDS were taken forward in the analysis (8312 unique probes, 6084 unique CDS, 49872 datapoints).

A Bayesian hierarchical model was fit to the array intensity data. This model treats growth and amplification (‘control’ and ‘treatment’ arms) and adherence to roots (‘treatment arm only’) as additive linear effects describing the relationship between the measured intensity for each probe *i* before (x_i_) and after (y_i_) each replicate experiment. In this model, parameters for the linear components were pooled either by the CDS from which the probes are derived (for gradients: β and δ, with corresponding index for the associated CDS *j*[*i*]), or the array used for that replicate (for offsets: α and γ, with corresponding index for the array/replicate *k*[*i*]). A binary 1/0 value (*t*_*i*_) was used to indicate whether a specific experiment did or did not include the spinach root adherence:

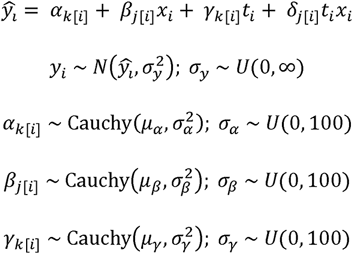

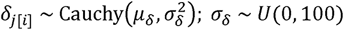

The model was fit using PyStan 2.12.0.0 under Python 3.6, with two chains each of 1000 iterations, to estimate parameter values: *α*_*k*[*i*]_ - the array-level offset due to growth for each replicate; *β*_*j*[*i*]_ - the C DS-level influence of the growth step on probe intensity; *γ*_*k*[*i*]_ - the array-level offset due to treatment/passage for each replicate; *δ*_*j*[*i*]_ - the C DS-level influence of treatment/passage on probe intensity; *μ*_*α*_, *μ*_*β*_, *μ*_*γ*_, *μ*_*δ*_ - the pooled distribution means for each of the four main equation parameters; *σ*_*α*_, *σ*_*β*_, *σ*_*γ*_, *σ*_*δ*_ - the scale values for the pooled distributions for each of the four main equation parameters; and *σ*_*y*_- the variance due to irreducible measurement error.

The C DS with index *j*[*i*] was considered to be associated with an advantageous effect on adherence (positive selection pressure) if the median estimated value of *δ*_*j*[*i*]_ was positive, and the corresponding 50% credibility interval did not include zero. A similar interpretation was used to infer an advantageous effect on *in vitro* growth/amplification from estimates of *β*_*j*[*i*]_ . Goodness of the model fit was estimated using 10-fold crossvalidation. The model is described in full in an interactive Jupyter notebook in Supplementary Information.

### 2.5 Molecular methods

All primers and plasmids are listed in Table S2. To identify BAC clones containing the *etp* operon, bacterial pools consisting of 48 clones of the library were screened by PCR for *etpD* and *etpO* genes using primers etpD.RT.F, etpD.RT.R, etpO.F, etpO.R. Individual clones in the pool were then screened using the same primers, identifying clone BAC2B5. BAC2B5 sequence was determined from primer walking near HindIII sites in pO157 with primers specific to the pVG1 vector. PCR products amplified using primer pairs BAC2B24F and pVG1; BAC2B5F and pVG1.R were Sanger sequenced. This confirmed the sequence from the BAC vector pVG1 to the upstream and downstream sequence at pO157 HindIII 87463. *E. coli* strain Sakai was cured of the pO157 plasmid by plasmid incompatibility as described by [28]. In short, Sakai was transformed with pBeloBAC11 which has the same incompatibility as pO157. Transformants were subcultured three times in LB+Cam to cure the pO157. Plasmid curing was confirmed by PCR for *toxB, hlyAB* and *etpO*. The pBeloBAC11 was cured by sub-culturing three times in LB without selection. Loss of pBeloBAC11 was confirmed by loss of Cam resistance and by PCR for the vector using primers T7 promoter and Cml_rev. The pO157-cured and WT strains were whole genome sequenced from a paired-end library to generate short-read (Illumina) sequences (ENA accessible number: ERS4383229 – accessible 30-Jun-2020), which were annotated using PROKKA [29] for Blastp [30] comparisons, using the reference Sakai sequence (BA000007.3) on the Galaxy platform [31]. A defined deletion in the *E. coli* O157:H7 isolate Sakai *etpD* gene (pO157p03) and *loc6* fimbrial locus (ECs1276-1280) was exchange as previously described [32, 14] using constructed vectors pAH005 (*loc6*) and pAH006 (*etpD*), respectively. The SakaiΔ*etpD* strain was cured for resistance to tetracycline by transforming the mutant with FLP recombinase expressing plasmid pCP20 [33]. Deletions were confirmed by PCR and Sanger sequencing, and for the Sakai Δ*etpD* strain by whole genome sequencing and BLASTn analysis to confirm loss of the CDS for pO157p03 locus. The promoterless *etpD* gene was PCR amplified (primers EtpD.Xba.pSE and EtpD.Hind.pSE) and cloned into the IPTG inducible plasmid pSE380 to create pAH007 and complement the mutation *in trans*. For the GFP transcriptional reporters, the 5’UTR of *etpC* and *etpD* was PCR amplified (primers EtpC.XbaI.F, EtpC.XbaI.R, pKC_EtpD.XbaF, pKC_EtpD.XbaR) and cloned into pKC026 using XbaI, creating the transcriptional fusions pAH008 (*etpC*) and pAH009 (*etpD*), respectively.

### 2.6 Bacterial adhesion assays on plant tissues

Adherence assays were performed as described in [15]. In short, plant tissues were washed and incubated in bacterial suspension (~1×10^7^ cfu/ml in sterile PBS; OD_600_ = 0.02) statically for two hours at 18 °C. Plant samples were vigorously washed 3 times in sterile PBS by mixing on a vortexer, weighed then homogenised with a sterile pestle and mortar. Samples were serially diluted and plated on MacConkey’s agar with appropriate antibiotics for bacterial counts. Measurements of *E. coli* Sakai wild type and *etpD* knockout, and Sakai Δ*etpD* transformed with the empty vector (pSE380) and *etpD* complement (pSE-*etpD*), were performed separately in batches of five biological replicates on independent leaf or root tissues as appropriate. Four batches were obtained for leaf tissue, and six for root tissue.

The bacterial recovery data (logCFU) was fit to a linear model describing additive non-interacting effects due to: *E. coli* Sakai adhesion (α); the modification of wild-type adhesion due to knockout of *etpD* (β); the introduction of empty pSE380 plasmid into the knockout background (γ); the effect of introducing pSE-*etpD* with respect to introduction of the empty vector in the knockout background (δ); and batch effects (ϕ_1..n_). The data were fit using PyStan 2.16.0.0 under Python 3.6, and the parameter estimates for β and δ and their 50% and 95% credibility intervals were used to infer the effects of knockout and complementation of *etpD*, respectively. These estimates represent the change in recovered bacterial counts as a result of the specific modification (loss or gain of *etpD*) with respect to the appropriate control. The model fit is described in full in a Jupyter notebook (https://widdowquinn.github.io/SI_Holmes_etal_2017/notebooks/04-etpD.html).

### 2.7 Bacterial adhesion to abiotic surfaces

Bacterial strains were cultured in LB at 37°C, 200rpm, for 16 hours then washed in fresh LB, RD MOPS glucose or RD MOPS glycerol. To assess initial attachment, the OD_600_ was adjusted to 0.5 for 2 hours incubation in the microtiter plate; for early biofilm formation, the OD_600_ was adjusted to 0.02 for 24 hours incubation. 200 μl was aliquoted in quadruplicate in an untreated 96 well plate (VWR, UK). The plate was incubated at 18°C statically before measuring adherent bacteria by Crystal Violet as described in [34].

### 2.8 Analysis of bacterial fluorescence *in vitro*

Gene expression was measured from *E. coli* Sakai transformed with pAH008 or pAH009 following growth for ~18 hours in LB medium + Chl at 37 °C, 200 rpm before diluting 1:100 into 15 ml RD MOPS medium supplemented with 0.2 % glucose or glycerol. Cultures were incubated statically at 18 °C and samples periodically removed and measured for cell density and GFP fluorescence. GFP fluorescence was measured in triplicate 200 μl volumes in a 96 well plate using GloMax plate reader (Promega). *E. coli* Sakai transformed with the vector control plasmid pKC026 was included as a control for background fluorescence. Fluorescence was plotted against OD_600_ and a quadratic line of best fit obtained. This was used to correct readings for background fluorescence. Corrected data was normalised to cell density (OD_600_) and values plotted using GraphPad Prism software for two experimental repeats.

### 2.9 Confocal microscopy

Fully expanded 4-week-old spinach leaves were infiltrated, by pressure injection using a 1 ml needleless syringe into the abaxial epidermis, with approx. 10^6^ cfu *E. coli* Sakai + pAH009 + *pmKate* and the plants maintained in an environmental cabinet until observed four days later. Two leaves on two individual plants were infiltrated per experiment and the experiment repeated on spinach plants propagated several weeks later. High inoculum levels ensured sufficient cells for observation since we have previously shown that *E. coli* Sakai is unable to proliferate in the apoplast of spinach and remains in a persistent state [17]. Leaf segments were infiltrated with sterile distilled water, to displace air from the apoplastic spaces between the spongy mesophyll cells, prior to mounting abaxial side up on microscope slides using double-sided tape. For spinach roots, 5 weekold spinach were grown under hydroponic culture as described, the 0.5× MS was removed and replaced with 10ml 0.5× MS inoculated with 10^8^ cfu bacteria. After four days in environmental cabinet conditions, the tub was flooded with sterile PBS to displace the perlite from the roots as non-invasively as possible. The leafy part of the plant was removed from the root by sterile scalpel cutting approximately 5mm below the cotyledon. After a further two washes in PBS, the root was mounted on a microscope slide, flooded with sterile PBS, and the coverslip held in place with double-sided tape.

Mounted plant tissue samples were observed using a Nikon A1R confocal laser scanning microscope mounted on a NiE upright microscope fitted with an NIR Apo 40× 0.8W water dipping lens and GaAsP detectors. Images represent false-coloured maximum intensity projections as indicated, produced using NIS-elements AR software. GFP (green) and chlorophyll (blue) were excited at 488 nm with the emissions at 500-530 nm and 663-737 nm respectively, and mKate (RFP) was excited at 561 nm with emission at 570-620 nm (magenta).

## 3 Results

### 3.1 Interaction screen using an *E. coli* isolate Sakai BAC clone library

To identify candidate gene loci for *E. coli* O157:H7 isolate Sakai (hereafter: *E. coli* Sakai) that conferred an advantage to spinach root tissue interactions, a Sakai BAC clone library was employed hosted in *E. coli* strain DH10B, which is derived from a K-12 strain and in our hands is a poor coloniser of plants [6]. A differential screen compared BAC clones inoculated with spinach roots to BAC clones treated similarly but in the absence of spinach roots. The BAC library was inoculated with freshly harvested spinach roots for two hours (insufficient time for bacterial proliferation) in four successive rounds to enrich for interactions. Loosely-attached and non-adherent bacteria were excluded between each round, so that the only strongly-adherent population were used for subsequent inoculation rounds, since these are most likely to be retained as ‘successful colonisers’. Each round resulted in successive reductions of the number of bacteria recovered from the roots as selectivity increased, with a 400-fold reduction between round 1 and 2 from 6 × 10^5^ cfu/ml to 1.6 × 10^3^ cfu/ml, which necessitated an amplification step after the second round to ensure that there were sufficient bacteria for subsequent selection rounds 3 and 4. An additional amplification step after round 4 ensured sufficient *g*DNA for hybridisation to the microarray. The no-plant negative control treatment did not include spinach root tissue, where the bacteria were inoculated into medium and suspended in PBS alone, to account for gene loci in the BAC clone library that may have conferred an advantage during the amplification steps between round 2 & 3 and after round 4. After four rounds of selection and enrichment, a total of 7.17 × 10^8^ cfu/ml of bacteria were recovered from the plant-treatment compared to 1.13 × 10^9^ cfu/ml of bacteria from the negative control treatment and taken forward for gene abundance analysis.

Gene abundance in pools of BAC clone *g*DNA was quantified on a DNA microarray before (i.e. input pools) and after selection (output pools), for both plant and no-plant treatments (dataset submitted to DiB DIB-S-20-00975). A Bayesian hierarchical model was fitted to the probe intensity data to estimate for each CDS in the *E. coli* DH10B and Sakai genomes a parameter representing the selection pressure due to inoculation on the plant. A CDS was considered to be under positive selective pressure (i.e. enriched) if its estimated value of this parameter was positive, and its 50% credibility interval did not include zero. This resulted in 115 CDS with a credible positive effect on adherence (Table S1).

### 3.2 Spinach root interactions enrich *E. coli* Sakai genes in six genomic regions (S-loops)

The 115 CDS that correlated with adherence to spinach tissue comprised seven contiguous regions of interest, of which 68 CDSs had existing functional annotation and 47 were annotated as hypothetical proteins (Table S1). Enriched genes were grouped by chromosome / plasmid location [10] and described in the context of the *E. coli* Sakai-specific S-loop designation [26]: S-loop 71; S-loop 72 / prophage SpLE1; S-loops 85 / prophage Sp9; S-loop 225; S-loop 231; and pO157 (Fig. 1).

**Figure 1.**
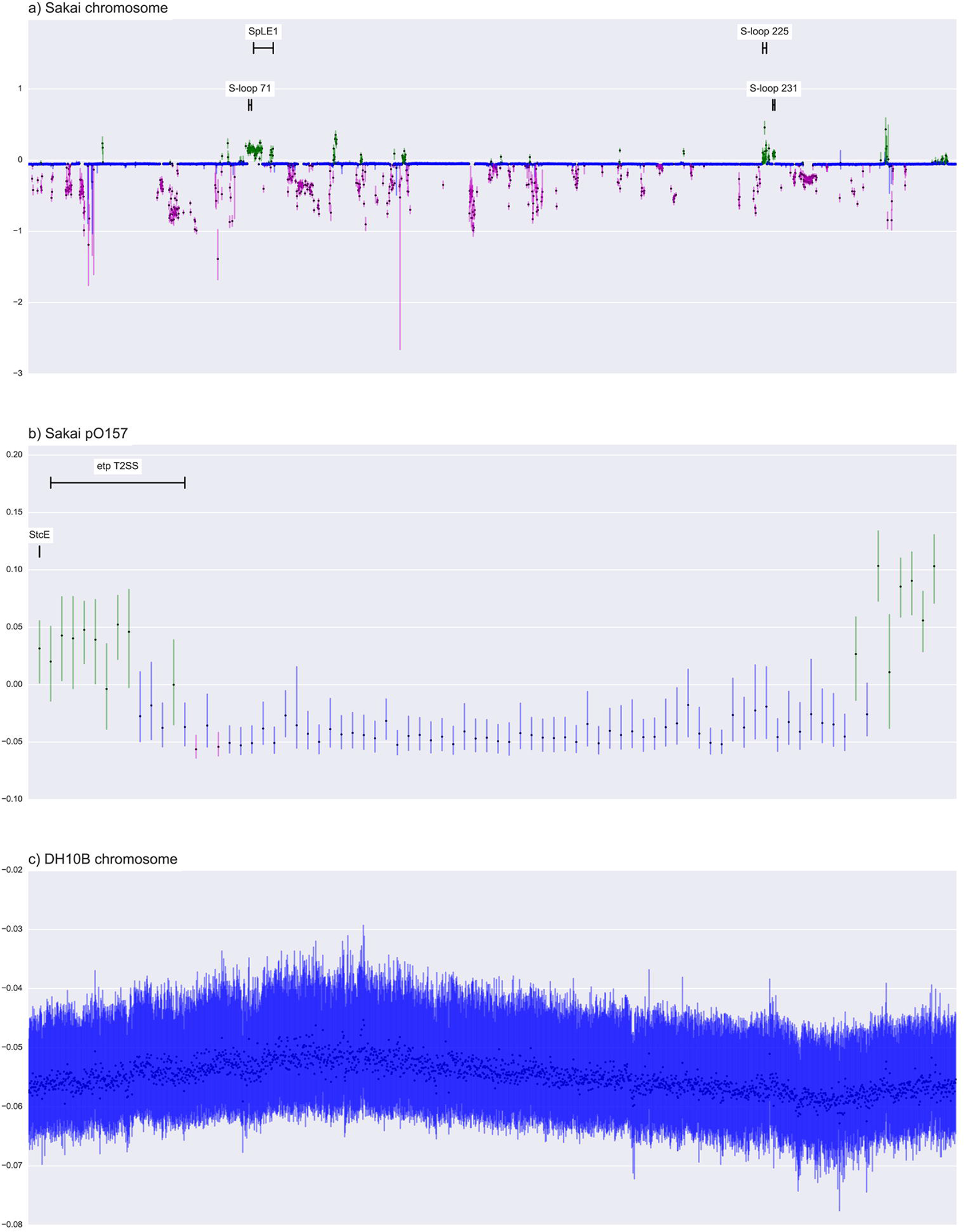
Regions in *E. coli* Sakai genome enriched by the adherence screen. Output from the model indicating estimated values of delta: the effect of treatment (passage) on retention of the introduced *E. coli* Sakai DNA in an *E. coli* DH10B background, for (A) Sakai chromosome DNA; (B) Sakai pO157 DNA; (C) the DH10B chromosome background. Estimated values are shown as black dots, and the 50 % credibility interval (CI) of this value as a vertical line. Where the 50 % CI does not include the median value for the dataset (assumed to represent a neutral response to passage), this may imply a selection response. Green CIs, where the median response is lower than the 50 % CI, are interpreted as positive selection pressure such that the gene is beneficial under passage. Magenta CIs, where the median response is greater than the 50 % CI, are interpreted as negative selection pressure such that the gene is deleterious under passage. Regions of the *E. coli* Sakai genome that are potentially under positive selection pressure include S-loop 71, S-loop 231, and S-loop 225; SpLE1, and the plasmid genes encoding the Etp type II secretion system, and StcE, as indicated. The DH10B chromosome genes show no evidence of positive or negative selection. Gene loci are listed in Table S1.

S-loop 71: a contiguous region in S-loop 71 was identified spanning 28 loci from ECs1272-ECs1296. This region is equivalent to the genomic island OI#47 in STEC isolate EDL933, which is conserved in STEC O157 serotypes [35], and includes the *loc6* fimbrial cluster, putative hemagglutinin/haemolysin-like proteins and fatty-acid synthesis genes.

S-loop 72: Sakai prophage like element 1 (SpLE1) in S-loop 72 encodes 111 open reading frames (ECs1299-ECs1409 [36]), of which 36 were enriched in interaction with spinach tissue, which we termed SpLE1 (partial). Enriched genes included those for urea degradation *ureA,B,EFG,* of which urease genes ECs1321-1327 were repressed in response to spinach root exudates [20]. Adhesion Iha and AidA, encoded by ECs1360 and ECs1396 respectively, are also present in SpLE1, but were not enriched in a contiguous region of 50 genes (ECs1349-1398).

S-loop 85: Prophage Sp9 in S-loop 85 includes a number of genes encoding non-LEE encoded (Nle) effectors (*nleA, nleH2, espO1-2* and *nleG* [37]). This region was enriched in a separate study investigating adherence to bovine primary tissue [23], and induction of *nleA* was induced in STEC (EDL933) in response to lettuce leaf lysates [19].

S-loop 225: Gene loci in S-loop 225 (ECs4325 – 4341) are associated with fatty acid biosynthesis and ECs4331 is annotated as a putative surfactin [26]. ECs4325-4340 were also induced in *E. coli* Sakai in the presence of spinach leaf lysates [20].

S-loop 231: Gene loci in S-loop 231 (ECs4379 – 4387) are associated with heme utilisation and transport and ECs4379 encodes a *chuS* heme oxygenase [38]. ECs4383/86/87 were induced in the presence of spinach root exudates [20] and locus Z4912 (ECs4381) was induced for STEC isolate EDL933 attached to radish sprouts [39].

pO157: pO157 p3,5,6, and 8 encode genes in the operon for a Type 2 secretion (T2SS) system. The T2SS of STEC has been reported to play a role in adherence to mammalian host tissues [40]. The pO157 has a role in biofilm formation, since a plasmid cured strain of *E. coli* Sakai was shown to have reduced EPS production and did not generate hyperadherent variants (Lim et al., 2010). Furthermore, the T2SS is an important virulence factor in many phytopathogens required for the secretion of plant wall degrading enzymes (reviewed in [41].

Analysis of the unclassified group (hypothetical genes) by InterProScan did not indicate any potential roles in adherence and none were selected for functional analysis: 18 had no predicted functional domains and six genes had a predicted transposase function (ECs1337-1340, Ecs3868-3869). Nine were included above: four *nle* effectors in prophage Sp9; urease gene ECs1321; fatty acid synthesis genes ECs4333 and 4335; and p79 and p81 from lipid operon *ecf*. Another four have domains of unknown function (DUF).

Oon basis of gene annotation and any reference in the published literature, we focused on two candidates that may have a function in adherence, as a key aspect of initial colonisation interactions: the *loc6* gene cluster from S-Loop 71 since fimbriae are well described adherence factors, and the T2SS genes on pO157, which are associated with biofilm formation. Therefore, the functional activity of *loc6* and the pO157-based T2SS was assessed with spinach tissue using a series of deletion mutants.

### 3.3 Functional characterisation of *loc6* fimbrial locus

A defined *loc6* (ECs1276-1280) deletion mutant was constructed in *E. coli* Sakai and its ability to interact with spinach roots compared to the WT parental strain. There was no difference between the numbers of the Loc6 fimbriae-deficient bacteria recovered compared to wild-type, following a two-hour incubation on spinach roots (Fig. 2). This suggested that the *loc6* fimbrial locus did not confer a direct advantage on spinach roots, and it is possible that genes elsewhere in the contiguous region were responsible for enrichment of the BAC clones (Table S1).

**Figure 2.**
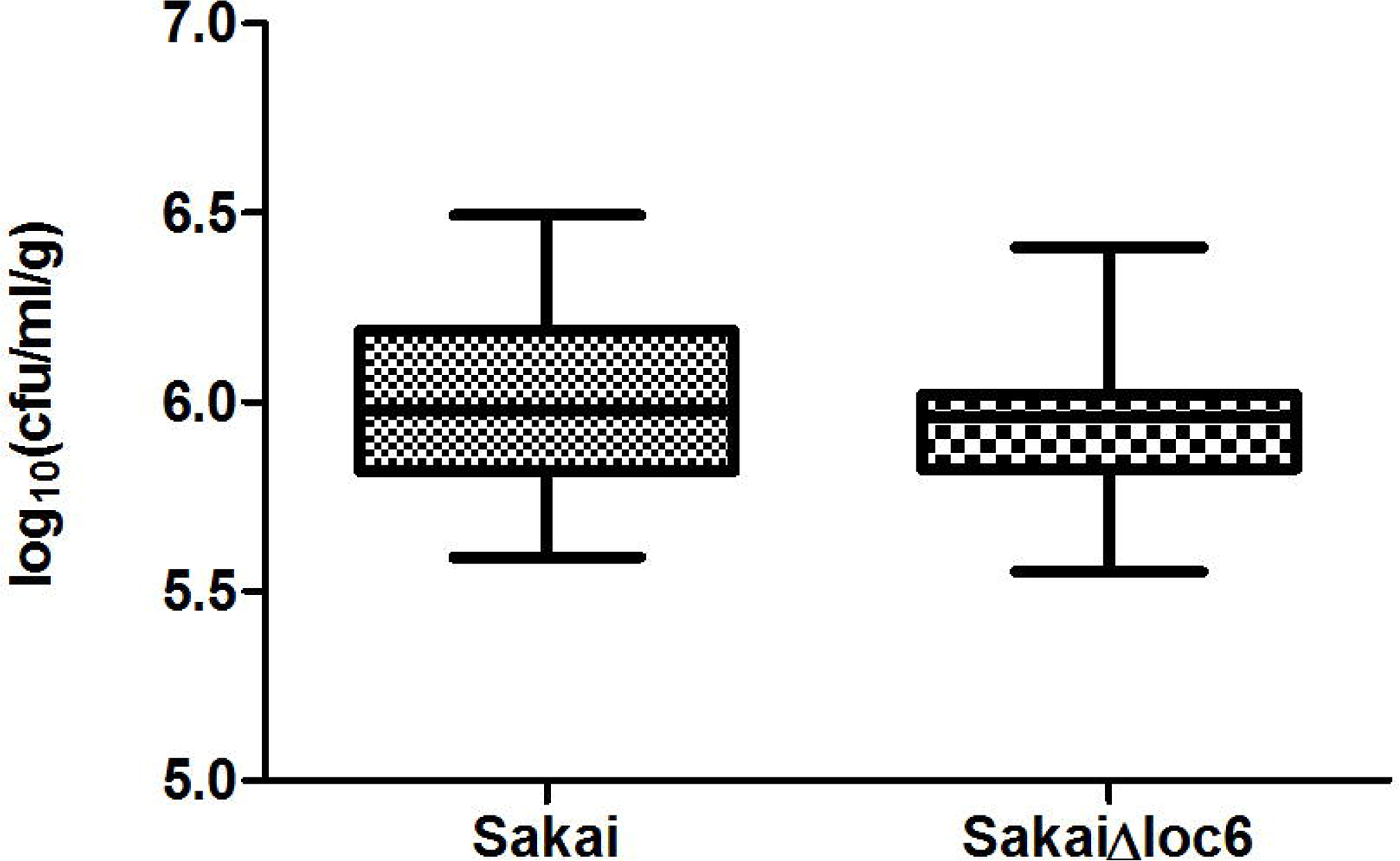
Assessment of *E. coli* Sakai Loc6 fimbriae in binding to spinach root tissue. *E. coli* Sakai or its isogenic *loc6* mutant recovered after a 2 hour adherence assay on spinach roots. The data from 3 independent experiments with 10 biological replicates for each bacterial strain are presented in box plots with the mean shown as a line in the interquartile ranges, and whiskers for maximum and minimum values. There was no statistically significant difference in the mean number of *E. coli* Sakai WT recovered compared to Δ*loc6* by Students *t* test (p=0.3268)

### 3.4 Functional characterisation of the pO157-encoded Type II secretion system

#### 3.4.1 A role for pO157 in spinach interactions

Candidate BAC clones containing TS22 genes (in *E. coli* DH10B background) were tested for their ability to interact with spinach root tissue compared to the empty BAC vector, pV41 (also transformed in DH10B). Clone BAC2B5, which encompasses the entire pO157 sequence, increased adherence to spinach roots significantly (p <0.05; students t test) compared to the pV41 vector-only control (Fig. 3A). A plant-dependent specificity of the pO157 BAC2B5 clone was determined by testing adherence to two non-plant surfaces. There was no significant difference in binding for clone BAC2B5 compared to the vector-only control on natural wool (a biotic surface mimicking root structures) (Fig. 3A; p=0.9864) or polystyrene (abiotic surface) (Crystal Violet (OD_590nm_) mean of BAC2B5: 0.0178 ± 0.0227; pVG1: 0.0236 ± 0.0303).

**Figure 3.**
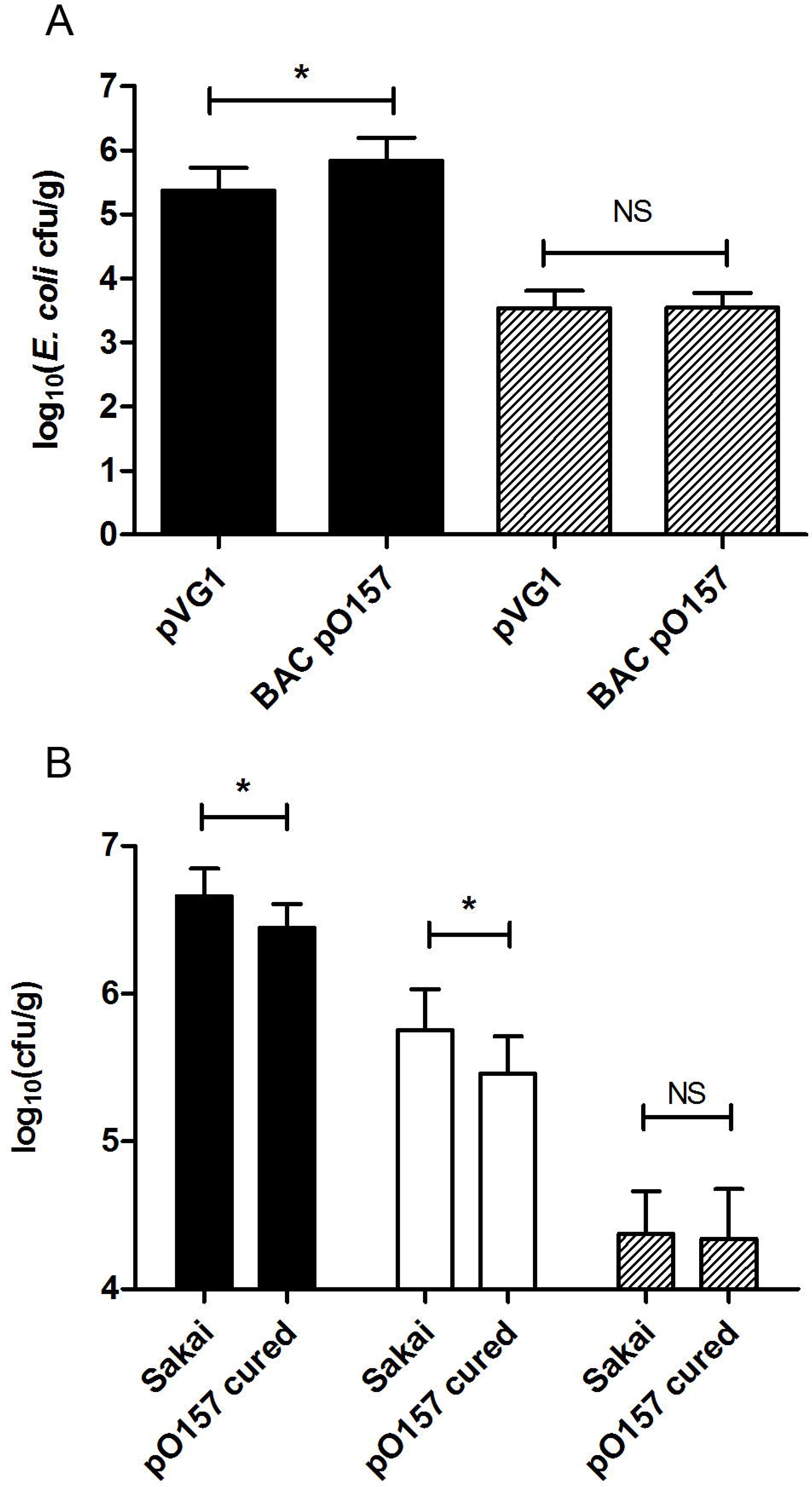
*E. coli* Sakai pO157 mediates interactions with spinach tissues. (A) *E. coli* DH_10_B transformed with BAC clone BAC2B5, containing pO157 sequence, or the empty BAC vector pV41 recovered from roots of hydroponics-grown spinach (filled bars) or natural wool (striped bars) and (B) *E. coli* Sakai WT or pO157-cured recovered from roots of compost-grown spinach (filled bars), leaves (open bars) or natural wool (striped bars). Data shown is the average from triplicate experiments each with five biological replicates. Statistical significance was calculated by students *t* test (* p<0.05, NS not significant).

A role for the pO157 plasmid in interactions with spinach was confirmed by removal of the pO157 plasmid from *E. coli* Sakai. Plasmid loss was confirmed by PCR for the pO157 specific genes *toxB, ehxA* and *etpO*, and from comparison of the whole-genome sequence and its isogenic parent (*E. coli* Sakai WT). All the annotated pO157 plasmid coding sequences were absent in the pO157-cured isolate except for two CDS associated with an IS element (IS629), while 100 % of the annotated chromosome and pOSAK1 plasmid CDS were present. The *E. coli* Sakai pO157-cured strain showed 99.996 % average nucleotide identity to the Sakai chromosome (GCA_000008865.2) (95.629 % alignment), with no or alignment to the pO157 plasmid, but partial coverage of pOSAK1 plasmid (100 % identity, 47.822 % alignment). Inoculation of *E. coli* Sakai pO157-cured with spinach plants significantly reduced the number of bacteria recovered from roots and leaves compared to its isogenic parent (Fig. 3B, black and white bars respectively). Binding to spinach tissue was not due to generic adherence to surfaces, since there was no significant difference between the number of *E. coli* Sakai pO157 mutant and its isogenic parent recovered from natural wool (Fig. 3B, wool grey bars).

##### 3.4.2 Analysis of a T2SS mutant in spinach interactions

To assess a role of the pO157-encoded T2SS in spinach binding, a defined knockout of the T2SS secretin protein, EtpD was constructed (*E. coli* Sakai Δ*etpD*). Whole genome sequencing confirmed the specific loss of the *etpD* CDS in its entirety, as designed. Average nucleotide identity between *E. coli* Sakai Δ*etpD* and the Sakai genome (GCA_000008865.2) showed 99.997 % identity to the chromosome (94.614 % alignment), and although short-read sequencing was performed, some contigs covered the plasmids, with 99.960 % identity to the pO157 plasmid (49.374 % alignment).

Adherence of the *etpD* mutant was compared to the isogenic parent to spinach roots derived from plants that were propagated in compost (Fig. 4B). Recovery of the *etpD* mutant (*E. coli* Sakai Δ*etpD*) was reduced by 0.32 logCFU (95% credibility interval −0.56:−0.09) compared to the control (*E. coli* Sakai WT), although adherence was not completely abrogated. Complementation of the *etpD* mutant with a plasmid-borne copy of *etpD* (*E. coli* Sakai Δ*etpD* + pAH007) under inducible control did not restore adherence to wild-type levels, relative to cells transformed with the empty vector control (*E. coli* Sakai WT + pSE380) also treated with the inducing agent, IPTG (Fig. 4). Substantial variation occurred between replicate plants and the average number of recovered bacteria with the empty vector (*E. coli* Sakai WT + pSE380) was greater than the *etpD* mutant without the plasmid (*E. coli* Sakai Δ*etpD*), indicative of an artefactual effect from the addition of IPTG. This was previously reported and suggests that IPTG may influence off-target genes that directly or indirectly alter adherence to plant tissue in *E. coli* Sakai [14].

**Figure 4.**
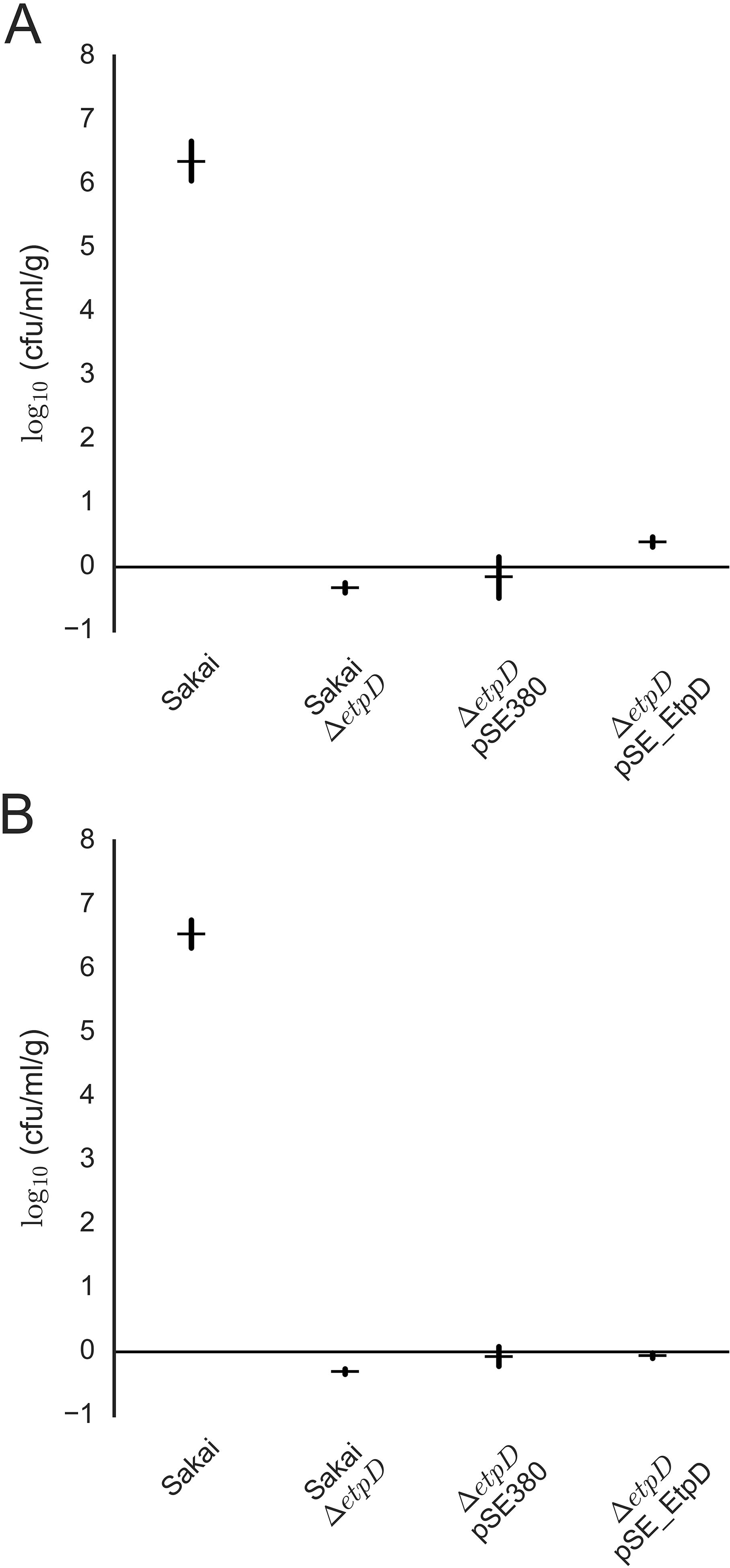
Modelling the impact of *E. coli* Sakai T2SS in interactions with spinach leaves and roots. Bacteria recovered from spinach plant tissue after 2 hour adherence assay. Regression coefficients (parameter estimates) obtained when fitting recovery data (CFU) from *E. coli* Sakai WT, Δ*etpD* and *etpD* mutant complemented with pSE380 or pAH007 (pSE_*etpD*) under IPTG-induction from leaves (A) or roots (B) to a linear model of additive effects, for each tissue. Sakai: expected recovery (logCFU) of wild-type *E. coli* Sakai; Sakai Δ*etpD*: expected (differential) effect on recovery of Δ*etpD* knockout with respect to wild-type Sakai; Δ*etpD* pSE380: expected (differential) effect on recovery of introducing the pSE380 into the knockout background; Δ*etpD* pSE_EtpD: expected (differential) effect on recovery of expressing EtpD, with respect to pSE380 alone. For each estimate, the marker represents the median value, and vertical lines represent the extent of the 50% credibility interval (50% of runs produce a value within this range).

The role in adherence for the T2SS was also tested on spinach leaf tissue to determine whether this function extended to other tissue sites. Recovery of the *etpD* mutant transformed with the empty vector (*E. coli* Sakai Δ*etpD* + pSE380) was enhanced with respect to the *etpD* mutant alone by 0.4 logCFU (95% credibility interval 0.15:0.63). Complementation of the *etpD* mutant using an inducible version of *etpD* cloned into single-copy plasmid (*E. coli* Sakai Δ*etpD* + pAH007) restored binding to 2.6-fold greater than the *etpD* mutant (*E. coli* Sakai Δ*etpD* + pSE380) (Fig. 4).

A plant-dependent specificity for *etpD* was confirmed by assessing binding to an abiotic surface (polystyrene), where there was no significant difference in attachment between *E. coli* Sakai Δ*etpD*, Sakai pO157-cured or *E. coli* Sakai WT, after either 2 hours (as measured by Crystal Violet, OD_590nm_ <0.050 ±0.025 SD) or after 24 hours, in 3 different media types.

##### 3.4.2 Expression of T2SS *in vitro*

The T2SS from *E. coli* Sakai is largely uncharacterised, both in terms of function and expression profile, with no data relating to plant-relevant environments. Therefore, expression was assessed from two independent plasmid-borne (multi-copy) transcriptional reporter fusions for *etpC*, the first gene of the operon, and for *etpD*, the outer membrane protein, since there is 211 nt between the stop codon of *etpC* and start codon of *etpD*, which includes putative transcriptional start sites (Fig. 5A). It appears that *etpD-K* are polycistronic since there is no apparent untranslated DNA between genes, and there is a predicted ribosome binding site upstream of *etpI*. The reporter fusions encompassed 508 nt and 257 nt upstream of the *etpC* and *etpD* start codons, respectively. Under *in vitro* conditions (defined medium at 18 °C), the maximum level of expression for both genes occurred in late exponential phase of growth (OD600 ~ 1), although there were marked differences in growth rates under the different carbon source regimes: *E. coli* Sakai reached this cell density in two days when grown with glucose, but needed six days with glycerol as a carbon source. The relative fluorescence was normalised to cell density to allow for comparison between the reporters, and GFP fluorescence from the *etpD-gfp+* reporter was five-to six-fold greater than the *etpC-gfp+* reporter (Fig. 5B). GFP fluorescence from both reporter constructs were three- to four-fold higher in RD-MOPS glycerol compared to that in RD-MOPS glucose; indicative of catabolite repression [42].

**Figure 5.**
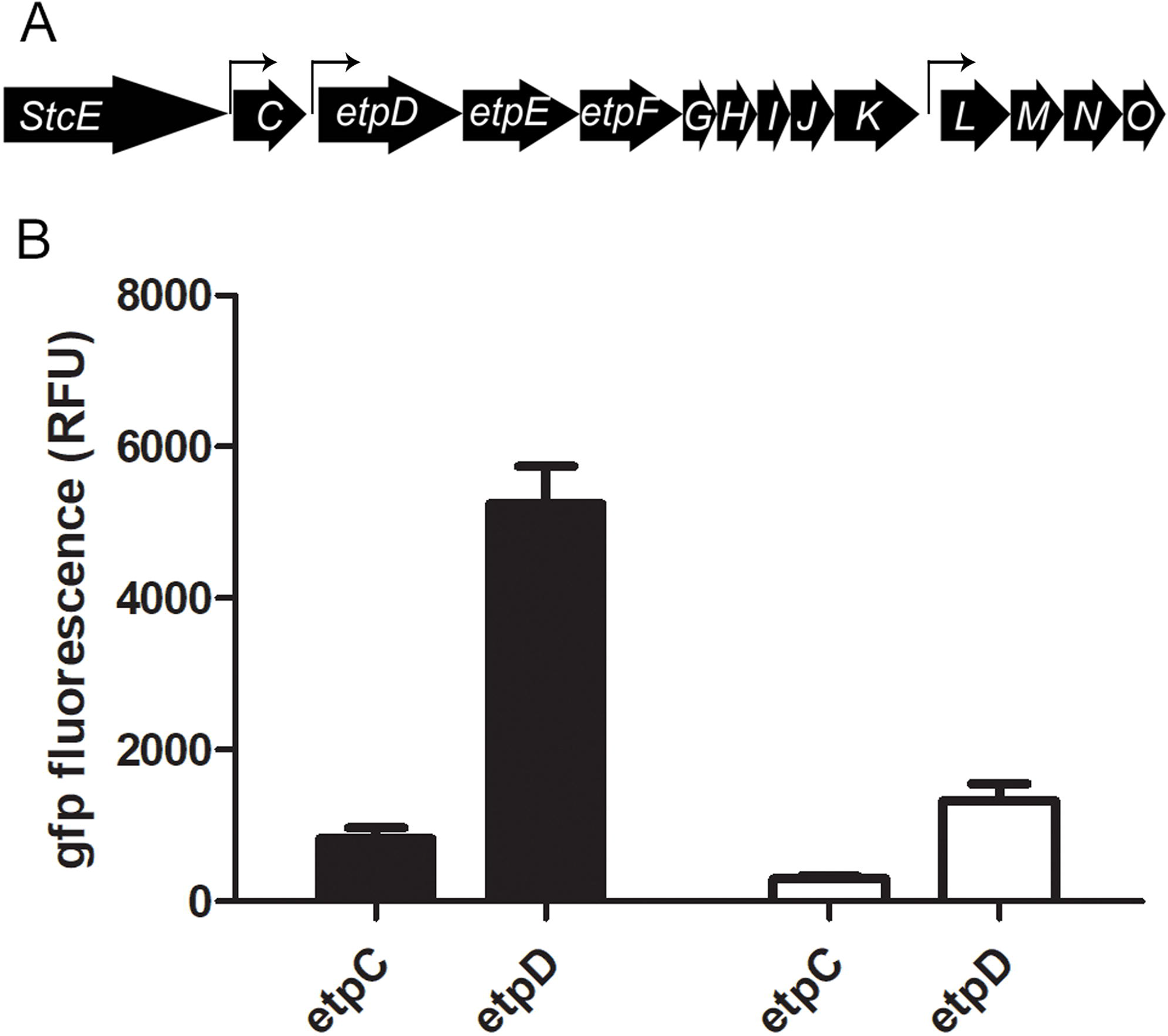
The *E. coli* Sakai *etp* T2SS operon and *in vitro* expression at 18 °C. Genetic organisation of the *etp* operon including the upstream metalloprotease gene *stcE* (A). GFP reporter activity for gene expression from the 5’UTR of *etpC* (508 bp) or *etpD* (211 bp) in *E. coli* Sakai, grown in RD MOPS medium supplemented with glucose (white) or glycerol (black). Expression values were corrected for background from the promoter-less reporter plasmid (pKC026) measured at the same optical density, and RFU normalised for cell density (OD_600_). Equivalent expression levels at late-exponential phase are provided (OD_600nm_ of 1) from two experimental repeats.

##### 3.4.4 Expression of the T2SS secretin gene, *etpD*, *in planta*

The transcriptional activity of the T2SS *etpD* secretin gene was assessed during *E. coli* Sakai colonisation of spinach roots or leaves, using the *etpD-gfp+* transcriptional reporter plasmid (pAH009). Repressive culture conditions for *etpD* expression (RD MOPS glucose: Fig. 5B white bars) were used to pre-culture the cells to observe *bone fide* expression, and *E. coli* Sakai + pAH009 were co-transformed with a constitutive RFP plasmid (p*mKate*) to aid location (Fig. 6). After four days, *E. coli* Sakai + pAH009 + p*mKate* were located along the surface of intact spinach root epidermal cells (Fig. 6A) or within an epidermal cell (Fig. 6B). Detection of GFP showed that *etpD* was expressed both on and inside spinach root cells, and expression was heterogenous, ranging from no GFP to very bright levels. Although the non-GFP expressers could have lost the reporter plasmid due to lack of selective pressure, detection of RFP from p*mKate* indicated maintenance of plasmids. *E. coli* Sakai located within the epidermal cell (Fig. 6Bi and ii) were apparently adherent to the plant cell wall (Fig. 6Biii), while others appeared to still be moving (since the plant tissue was live and unfixed during imaging) (Fig. S1A, arrow). *E. coli* Sakai co-transformed with p*mKate* and a constitutive GFP reporter (*pgyrA-gfp*) showed that the experimental conditions did not impact GFP detection and resulted in a similar pattern of colonisation, with apparently adherent cells (Fig. S1A, circle), indicating that harbouring two plasmids did not incur detrimental effects on isolate Sakai colonisation. As expected, there was no GFP observed from *E. coli* Sakai co-transformed with p*mKate* and the no-promoter pKC026 plasmid vector control (Fig. S1B).

**Figure 6.**
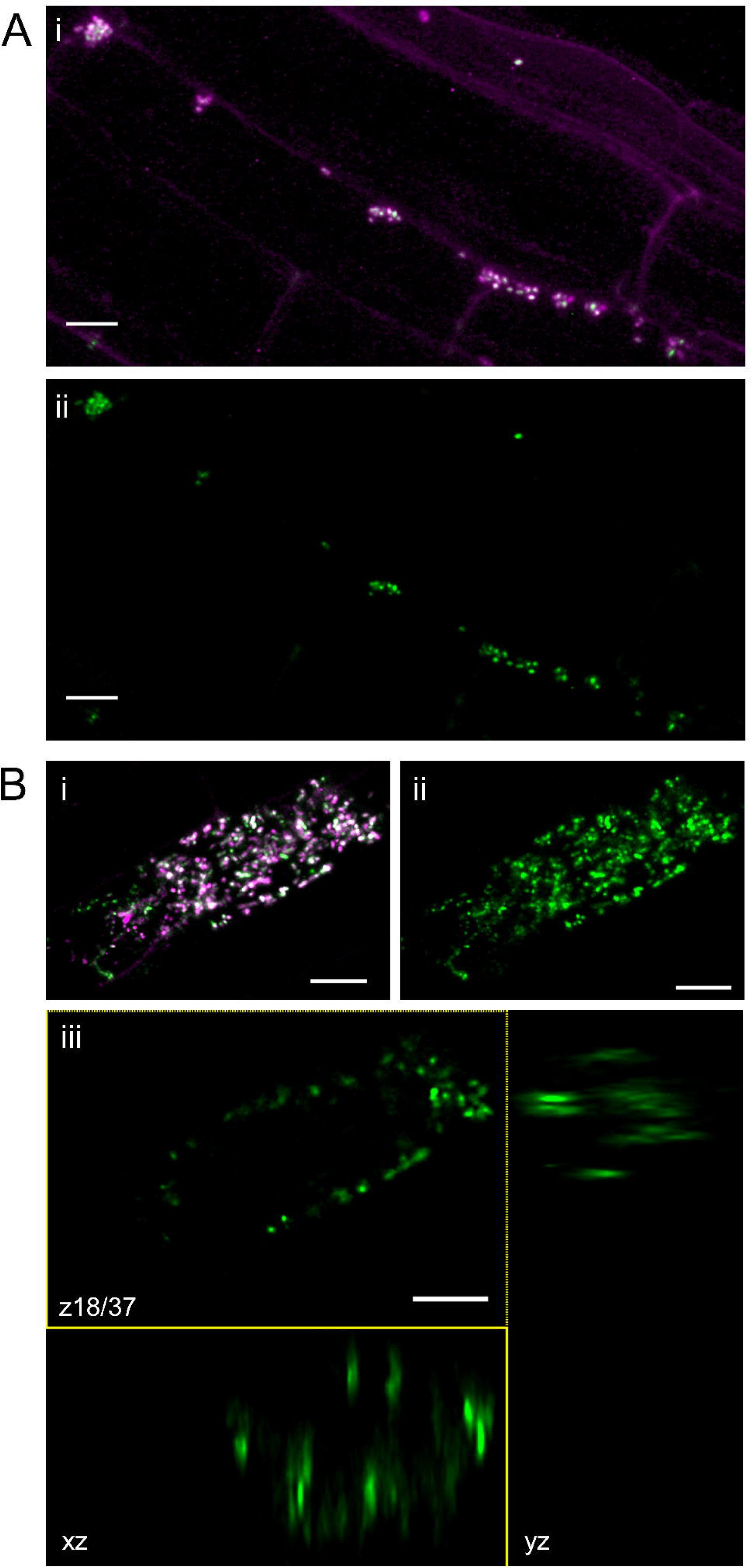
Expression of *E. coli* Sakai *etpD* during root colonisation. Spinach roots inoculated with 10^8^ cfu of *E. coli* Sakai co-transformed with pmKate and pAH009 (*etpD-gfp+*) were imaged by confocal microscopy after 4 days. *E. coli* Sakai were located along (A) or within (B) root epidermal cells with some *E. coli* Sakai attached to the cell wall within an epidermal cell (Biii). Maximum intensity projections (A, Bi-ii) of root epidermal cells with the merged image (Ai, Bi) or green channel (Aii, Bii-iii). GFP expression in green and RFP expression in magenta; root cell wall autofluorescence is also detected in the magenta channel (Ai). Scale bars are 10 μm. The panel of images are representative of four independent experiments from individual plants.

Expression of *etpD* was also shown for endophytic *E. coli* Sakai +pAH009 + p*mKate* located within the apoplast of spinach leaves (Fig. 7), from individual cells attached to spongy mesophyll cells (Fig. 7A) or adjacent to the cell wall (Fig. 7B), and in small chains of cells (Fig. 7C). In contrast, no GFP was observed from *E. coli* Sakai transformed with empty vector control (pKC026) (Fig. 7D).

**Figure 7.**
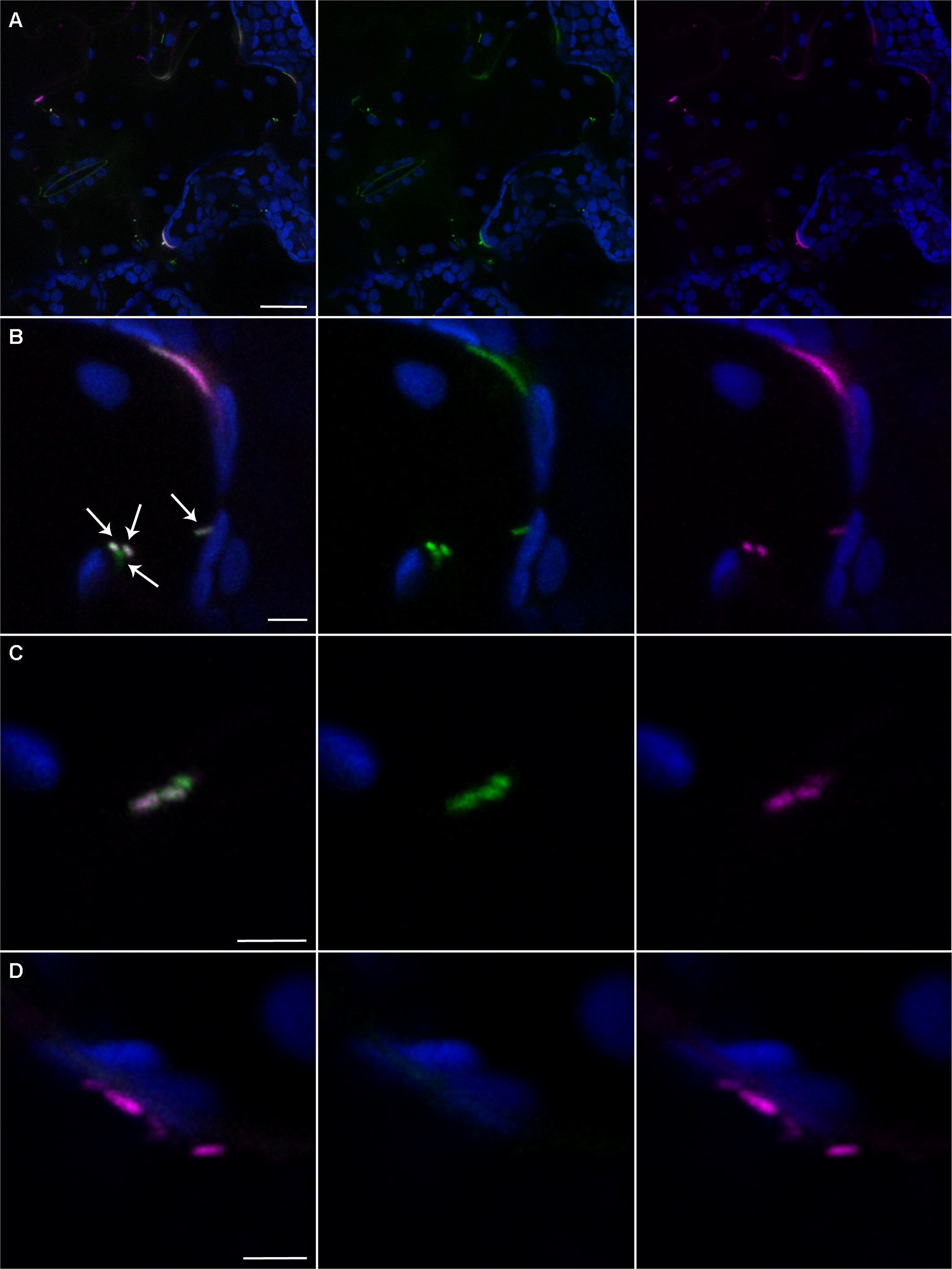
Expression of *E. coli* Sakai *etpD* in spinach leaves. Spinach leaves infiltrated with *E. coli* Sakai co-transformed with pmKate (constitutive expression of RFP) and pAH009 (*etpD-gfp+*) (A-C) or promoterless *gfp+* vector pKC026 (D) were imaged by confocal microscopy after four days. Chloroplast autofluorescence is false coloured blue in the images; GFP expression in green and RFP expression in magenta. Three sets of parallel panels show the maximum intensity projection of abaxial epidermal and mesophyll cells with the merged image (left), green channel (centre) and red channel (right). The panel of images are representative of two independent experiments from individual plants. Scale bars are 25 μm(A) or 5 μm (B-D). Examples of co-expression of *etpD-gfp* and *rfp* are indicated by white arrows (B).

### 4 Discussion

The main aim of this study was to identify novel STEC genes that mediate early interactions with fresh produce plant hosts. A high-throughput positive selection approach was used, where a BAC library of *E. coli* Sakai genomic fragment clones was screened for interactions to spinach roots. Spinach has been linked with high profile outbreaks of STEC, and although plant roots are not consumed they represent the preferred site of colonisation of *E. coli* Sakai. The screen enriched for the equivalent of 2 % of the *E. coli* Sakai genome, which is in-line with other studies using alternative approaches, e.g. a whole transcriptome study of *E. coli* Sakai identified two or six ‘adherence’ genes following inoculation with lettuce plants for one hour or two days, respectively (Linden et al., 2016). Several of the enriched gene loci were previously reported for STEC interaction with plant tissue, validating both the screen and their potential plant-associated functional role.

Adherence is a key step in early interactions with host tissue and STEC fimbrial adhesins that mediate specific binding to plant cell wall components include *E. coli* common pilus (ECP) and Yad fimbriae [14, 43] and non-specific interactions via flagella [15]. Potential candidates enriched in the screen may be involved in non-adherence functions, such as response to PAMP perception by the host, Nle effectors since NleA is known to play a role in disrupting secretory pathways [44, 45] or modulating host cytoskeleton (EspO1-2) [46] in animal hosts. Metabolic processes are also key for colonisation, which may explain enrichment of siderophore, ChuS.

One of the enriched loci selected for functional assessment on the basis of potential adherence included a chaperone-usher fimbrial gene cluster, termed Loc6 [11], was previously shown to be induced in STEC isolate EDL933 (gene Z1536) 30 minutes after exposure to lettuce leaf lysates [19]. In a separate study, the gene encoding the outer membrane protein (ECs1277) was induced in *E. coli* Sakai in response to a temperature reduction, to 14 °C [47]. However, the absence of any positive interaction with spinach root tissue indicated either no functional role or a subtle effect on binding. Alternative genes in the contiguous region identified by the BAC screen that may have contributed to interactions include a two-partner secretion (TPS) system termed *otpAB* (ECs1282-1283), which was characterised in STEC isolate EDL933 [48] and shares 100% sequence identity with *E. coli* Sakai. Although OtpA and OtpB apparently constitute a genuine TPS system in this isolate, the gene sequences did not genetically cluster with either of the two major subtypes of characterised two-partner secretion systems, haemolysins or adhesins [48]. Therefore, the authors postulated that the function of *otpA* could be accessory to that of the upstream fimbrial locus (*loc6*), which suggests that there may be a linked function between the gene clusters.

The second enriched candidate region selected for functional analysis was the T2SS encoded on the *E. coli* Sakai plasmid, pO157. The pO157 plasmid is ~ 93 Kb and also encodes virulence factors such as haemolysin genes, a catalase, a serine protease and a toxin gene [49]. The T2SS is widespread but not ubiquitous in bacteria and has been reported for bacteria from a range of hosts and environmental habitats [50]. In the related phytopathogen *Pectobacterium atrosepticum*, the T2SS (termed the Out system) bears structural and evolutionary similarity to the conjugative T4 pilus, and the gene cluster organisation tends to be labelled with gene ‘C’ at the beginning and gene ‘O’ at the end of the cluster. It is often termed the general secretory pathway (*gsp*), but in *E. coli* it is termed the <u>E</u>HEC <u>t</u>ype II <u>p</u>athway (*etp*) [51]. EtpD is orthologous to the secretin protein, ‘D’ that forms a channel across the outer member, while EtpC is homologous to the ‘C’ protein that spans the innermembrane as an anchoring protein [50]. Outside the *Escherichia* genus, EtpC has lower levels of homology to other species T2SS than EtpD [51], but does retain the functional domain of the superfamily of PulC proteins [52].

Absence of the pO157 plasmid reduced the number of bacteria recovered from spinach tissue, which appeared to be dependent on the EtpD secretin protein. Gene expression analysis supports a role for the T2SS *in planta*. The T2SS was shown to be responsive to incubation with plant tissue, with induction of *etpC* in response to spinach leaf lysates and spinach root exudates, and *etpD* induced in response to spinach root exudates [20]. Here, we show that expression occurs at plant-relevant temperatures (18 °C), and that both *etpC* and *etpD* expression was induced in the presence of glycerol but not glucose. Our data also supports independent promoter activity for both genes, albeit to differing levels. It is notable that the *etp* gene cluster for *E. coli* Sakai is encoded on the pO157 plasmid, whereas in other *E. coli* pathotypes the genes are chromosomal, indicative of recent recombination events, which could influence regulation in a background-dependent manner. A role for the STEC T2SS in colonisation of plant hosts is supported by data that shows the *etp* genes were upregulated in spinach outbreak STEC isolate TW14359 compared to *E. coli* Sakai upon adherence to mammalian MAC cells *in vitro* [53]. However, expression of the T2SS was not a pre-requisite for colonisation of bovine GI tract [24, 54] or gnotobiotic piglet intestines [55], indicating a degree of specificity in its function.

Whether or not the TS22 interacts directly with plant tissue, or indirectly via a T2-secreted protein, is not yet clear. Functional analysis of the T2SS in STEC isolate EDL933 showed that it is required for secretion of StcE (TagA), a metalloprotease that cleaves a C1-esterase inhibitor (C1-INH) [56], glycoprotein 340 (gp340) and mucin7 [57]. A role for the T2SS binding to mammalian tissue was demonstrated with Hep-2 cells [57], HeLa cells and in colonisation of the rabbit intestine [58]. Beyond that there is little available information on the STEC T2SS.

#### 4.1 Conclusion

High-throughput screening of the *E. coli* Sakai genome, using a BAC clone library, has enabled identification of a novel role for the T2SS of this foodborne pathogen. We have shown that it is expressed under relevant plant-host conditions and its presence enhances the short-term interactions of *E. coli* Sakai with plant hosts. Given the widespread nature of the T2SS, and a proven plant-colonisation role for T2SS of phytopathogens, it is perhaps not surprising that the STEC T2SS can mediate plant colonisation interactions.

## 5 Acknowledgments

The work was supported by a BBSRC grant to NJH, AH & LP (BB/I014179/1); Scottish Government Strategic funding to NJH, PH and JM; and BBSRC Institute grant funding to DLG and SPM (BB/J004227/1). Thanks to Steve Whisson for the gift of pBeloBAC11, Kathryn Wright for assistance with confocal microscopy and to Jacqueline Marshall and Marta Lis for technical assistance.

## 6 CRediT author statement

**Ashleigh Holmes**: Investigation, Formal analysis, Writing - Original Draft; **Leighton Pritchard**: Investigation, Formal analysis, Writing - Original Draft; **Peter Hedley**: Validation, Data Curation; **Jenny Morris**: Investigation; **Sean P. McAteer**: Resources; **David L. Gally**: Resources, Funding acquisition; **Nicola J. Holden**: Conceptualization, Writing - Review & Editing, Supervision, Funding acquisition

## 8 Tables

**Table S1** Description of gene loci enriched by the adherence screen, indicating S-loop and genomic location; output data from the enrichment analysis; gene annotation and description.

**Table S2** Description of plasmid and primers used in the study.

## 9 Figure Legends

**Supplementary Figure 1** Confocal microscopy controls of spinach root colonisation

Spinach roots inoculated with 10^8^ cfu of *E. coli* Sakai co-transformed with pmKate and *pgyrA-gfp+* (A) or empty vector pKC026 (B) were imaged by confocal microscopy after 4 days. Maximum intensity projections (left column A and B) of spinach root epidermal cells colonised by *E. coli* Sakai, within (A) or on the surface (B) of the cell. Volume projection of spinach root epidermal cell (A right column) showing *E. coli* Sakai colonisation within the epidermal cell with bacteria attached to the plant cell wall (circled). *E. coli* Sakai may also have been moving during image acquisition (arrow). GFP expression is coloured green and RFP expression in magenta. Scale bars are 10 μm. The panel of images are representative of four independent experiments from individual plants.

## Notes

### Competing Interest Statement

The authors have declared no competing interest.

