## Supplementary figures and images for "A high-throughput genomic screen identifies a role for the plasmid-borne Type II secretion system of *Escherichia* coli O157:H7 (Sakai) in plant-microbe interactions"

### Supplemental Figure 1

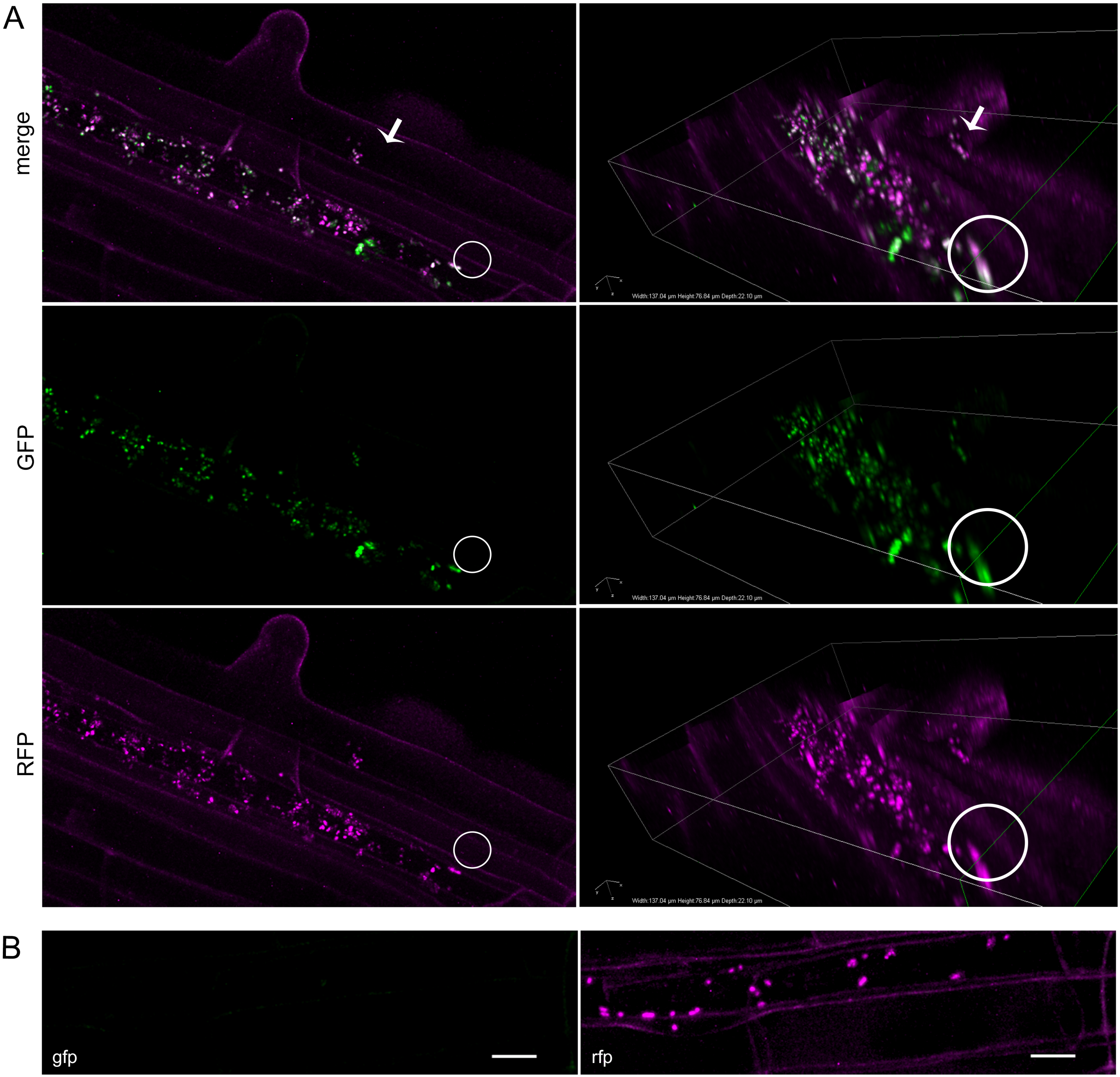
